# Effect of light intensity and nutritional value of food resources on flight response of adult parasitoid, *Cotesia plutellae* (Kurdjumov) (Hymenoptera: Braconidae)

**DOI:** 10.1101/444224

**Authors:** Kapinder, Tarkeshwar, Ashok Kumar Singh

## Abstract

*Cotesia plutellae* (Kurdjumov) (Hymenoptera: Braconidae) is the major larval parasitoid of *Plutella xylostella* (L) (Lepidoptera: Plutellidae), which is a serious pest of cruciferous plants throughout the world. We evaluated the influence of light intensities and feeding conditions on the vertical angle of flight in freshly emerged wasps in a cylinder having diameter 15cm and height 30cm. Light intensity was found to directly affects the flight activity. Increase in light intensity causes increase in vertical flight of the female wasps. However, Increase in light intensity did not influence the inclination of vertical flight in males. Feeding condition was also found to affect the vertical flight of the wasps. Honey odour, from below the flight chamber, arrested the flight of unfed or sucrose fed wasps. However, flight of honey fed wasps was not affected by honey odour. Male flight response was also influenced by feeding condition and light intensity but the response was not as higher as shown by females. The present study is useful for selecting suitable food prior to inundative release of parasitoid in the field at suitable time period of the day.

## Introduction

Among insects, parasitoid wasps are ecologically very important group and play a decisive role in an ecosystem by controlling their host’s abundance. The efficiency of using natural enemies to control pest under field conditions largely depends on their mobility (Mills and Heimpel 2018) and more specifically on their capacity to quickly locate pest infestation as well as to locate food resource like floral nectar or extra floral nectaries (Wanner et al. 2006; Yu et al. 2009). The dispersal capacity and propensity, without assistance, vary among different parasitoid species such as *C. glomerara* (Wanner et al. 2006) and Microplitis mediator (Yu et al. 2009) which depends on food prior to flight. For many natural enemies, for example, parasitoids mobility is directly related to flight aptitude which is determined by capacity and inclination of a species engaged in flight. There are several biotic factors such as body size, age, sex (Fahrner et al. 2014; Alves et al. 2015; Gaudon et al. 2016; Gaudon et al. 2018) and abiotic factors, like temperature (Rousse et al 2009; Jerbi-Elayed et al. 2015; Gaudon et al. 2016), photoperiod (Alves et al. 2015) and humidity (Rousse et al. 2009; Keppner and Jarau 2016) plays major role in determining the host searching and flight activity of insect parasitoids. Out of these, light intensity and feeding status are two important factors which directly affect the flight behaviour of insects.

Response to change in light intensity is important in habitat orientation. Andrewartha and Birch (1954) suggested that an organism may become adapted to respond to a gradient in intensity or to the quality of light and light can thus act as a “token stimulus” leading the animal to a place where temperature or moisture is favourable, to a place where there is an abundance of food, etc.

In the field, adult parasitoids are observed visiting flowering plants to feed on floral or extra-floral nectar (Jervis et al. 1993; Idris and Grafius 1995) to access carbohydrates and water which is essential for parasitoid maintenance, especially for increased flight capacity (Wackers 2005). This is true for *T. planipennisi* (Fahrner et al. 2014), *C. glomerata* (Wanner et al. 2006), and several other parasitoid species. The energy obtained through adult feeding on supplemental sugar sources may also be necessary for sustaining flight capacity in parasitoids because flight in insects is a highly energy-demanding behaviour with metabolic rates during flight increasing 50–100- fold compared with metabolism at rest (Beenakkers et al. 1984; Chapman, 1998; Hoferer et al. 2000, Fahrner et al. 2014). In most agro-ecosystems where parasitoids play a crucial role in controlling insect pests’ population, the two vital resources (i.e. host insect and food resource) for parasitoids are often not present in the same place. So, parasitoids must divide their time between host foraging and food foraging (Jervis et al. 1993; Eijs et al. 1998; Lewis et al. 1998). The time spent foraging for food detracts from the time available for host searching. As a result, food foraging can be viewed as a trade-off between an investment in future reproduction and immediate fitness (Sirot and Bernstein 1996).

To overcome this, adult parasitoids should be successful flier which can move frequently by flight between host and food containing areas to gain maximum reproductive success (Lewis et al. 1998). Thus, the availability and the quality of floral nectar sources may determine, to a large degree, the flight capacity of parasitoids and the range they can search for hosts, thereby influencing the efficacy of parasitoids as biological control agents. However, little is known about the influence of feeding carbohydrate rich food on flight capacity, and hence dispersal efficiency as well as flight inclination in parasitoids, due to the general lack of empirical studies. So, the first part of this study explains the influence of light intensity on the flight response of both the sexes of adult *C. plutellae* in cylindrical flight chamber. Secondly, influence of feeding status on the flight activity in adult *C. plutellae* was also evaluated in this paper.

## Materials and methods

### Insects culture

Larvae of diamondback moth (DBM), *Plutella xylostella,* parasitized by adult *Cotesia plutellae* were collected from cauliflower plants near Delhi (India) and reared on its leaves in the laboratory at 25 ± 1°C under 65 ± 5% R.H. and 14L: 10D photoperiod. The pupae formed by the parasitoid larvae emerging from the DBM were placed in separate jars for emergence. The emerging wasps were transferred to a clear acrylic ventilated chamber (20 x 20 x 20 cm) and given 50 % honey in water as food. These wasps were allowed to mate, and oviposit on late second instar DBM larvae which were transferred to clean jars and reared on cauliflower leaves in the absence of honey.

For each test, required number of parasitoid cocoons were drawn from the culture and kept one each in a glass vial (60 mm long, 15 mm dia). Virgin, water satiated and food-deprived wasps were used for various tests within 4 – 8 h of their emergence. All tests were carried out in a room with exhaust facility and maintaining 25±1°C and 65±5% R.H.

### Methods to study flight response

The influence of light intensities and feeding status on the vertical angle of flight in freshly emerged naïve male and female *C. plutellae* was evaluated in flight chamber under laboratory condition. The experiment was carried out in a cylinder having diameter 15cm and height 30 cm. The cylinder was divided into three zones lower, middle and upper zones. The inner wall was made sticky by an odour less grease and top was covered by sticky transparent lid. The whole setup was covered by black paper except at top from where light was allowed to come. The chamber was illuminated from above by two 20 W fluorescent tube lights (60 cm each) at a height of 20 cm from the horizontal arm. The intensity of light was controlled by the dimmerstat.

### Effect of light intensity

The flight angle of inclination was measured for both males and females at different light intensity in cylindrical flight chamber. The intensity of light used were 0 lux, 10 lux, 100 lux, 500 lux and 1000 lux. A group of five freshly emerged wasps were released in the middle of the base of flight chamber. The chamber was undisturbed for one hour. After that, the upper lid was open and the wasp’s stuck to the different zones of the cylinder were counted. The experiment was repeated five times of 50 insects arranged in 10 insects of each replicate.

### Effect of feeding

Freshly emerged wasps were allowed to feed on honey solution (50%) for fifteen minutes. After feeding, wasps were kept in the vials. The 10 ml of 50 % honey solution was soaked on whatman filter paper no. 1 in petridish, of 15 cm diameter,. The petridish was kept at the base of flight chamber and covered with nylon net of mesh size 12 x 12 mesh /cm so that insect could not touch the honey and could only perceive the volatile. Thirty minutes after feeding, a groups of five insects were released in the base of the chamber. The chamber was undisturbed for one hour. After one hour, upper lid was removed and the insects stuck to different zones of the cylinder were counted. The light intensity of 100 lux was provided from the top of the chamber. Similarly, insects were fed on 20 % sucrose solution and flight was tested in the presence of honey volatile provided from below the chamber. The experiment was repeated five times and each replicate consists of 10 insects.

### Statystical analysis

One way ANOVA was performed for flight activity at different light intensity and feeding condition between male and female. Student t-test was performed to compare flight ability of wasp in different light and feeding condition. All the statistical analyses were carried out using the computer program SigmaStat 2.0.

## Results

### Flight at Different Intensity of Light

Activity of *C. plutellae* was evaluated in terms of vertical flight in the flight chamber as described in materials and methods. With increasing light intensity, a significant increase (P<0.05) in flight response of female wasps to the upper zone was observed. Whereas, significant decrease (P<0.05) in the flight response was observed at lower zone. At 0 lux, female showed poor flight response, either they did not take off or they remained stuck to lower zone of the chamber. The percentage of female wasps that stuck to the lower zone (78%) was significantly higher (P<0.05) than middle and upper zone (12% and 10% respectively) (Fig. 1). However, there was no significant difference between middle and upper zone.

**Fig.1.**
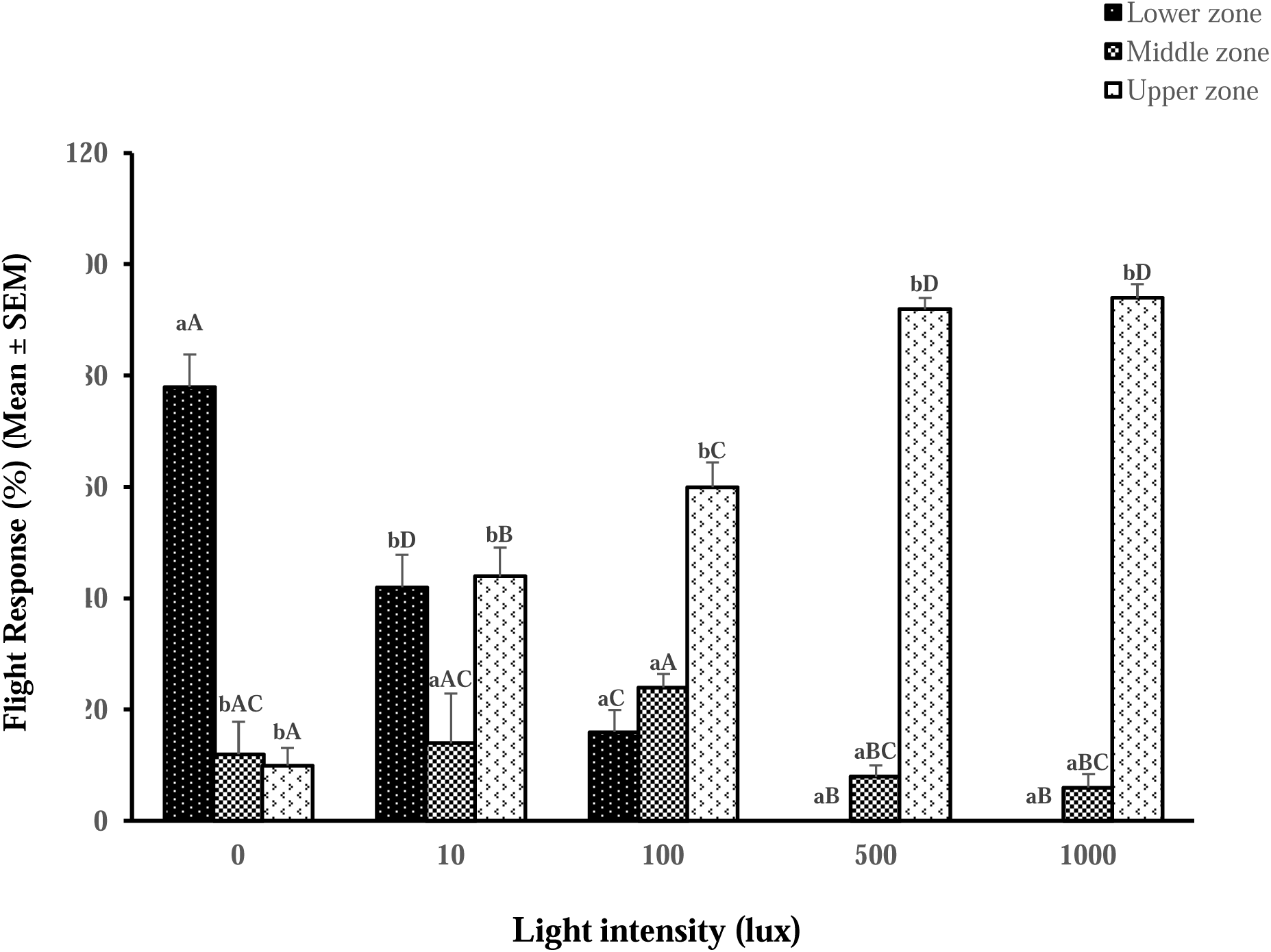
Flight response of female *C.plutellae* wasps at different light intensities in a cylindrical flight chamber. (Bars with different lower case letters differ significantly (p<0.05) on different zone of same light intensity. Bars with different upper case letters differ significantly (p<0.05) between different light intensity at particular zone)

At 10 lux, percentage of female wasps that stuck to the lower and upper zone was statistically similar (42% and 44% respectively) (P>0.05) whereas, the percentage of female sticking to the middle zone (14%) was significantly lower (P<0.05) than other two zones of the chamber. At 100 lux, vertical flight of female wasps increased and significantly (P<0.05) higher number of these wasps stuck to the upper zone (60%) as compare to lower (16%) and middle zone (24%). Whereas, sticking of wasps to the lower and middle zone did not differ significantly (P>0.05) (Fig. 1). Further increase in flight activity was observed at 500 lux. Significantly higher percentage of female wasps was observed sticking to the upper zone (92%) as compared to lower (0%) and middle zone (8%) (Fig. 1). Whereas, no difference was found between lower and middle zone of the chamber. Similar trend was observed at 1000 lux, where the percentage of female wasps sticking to the upper zone (94%) was significantly higher (p<0.05) than at the other two zones.

The vertical flight of female *C. plutellae* wasps was also compared in between various light intensities at a particular zone. At lower zone, the percentage of female found sticking was highest at 0 lux intensity (78%) and none was found at 500 lux and 1000 lux intensity. There was significant difference (P<0.05) in the percentage of female wasps found sticking to the lower zone between light intensities 0, 10, 100 and 500 lux. However, the difference was not significant between 500 lux and 1000 lux. It was found that the female wasps that stuck to lower zone, decreased significantly, as the light intensity was increase from 0 lux to 500 lux. However, at upper zone, the percentage of the female wasps stuck was directly proportional to the light intensity.

The flight of male wasps was also influenced by light intensity and the trend was similar to that of female wasp (Fig. 2). However, the flight response in them was weaker than female wasps. At 0 lux, significantly higher (P<0.05) percentage of male wasps stuck to the lower zone (90%). Whereas, the percentage of wasps that stuck to the middle (8%) and upper zone (2%) did not differ significantly (P>0.05) with each other. At 10 lux, only 24% of the male were stuck to the upper zone while 58% remained to lower zone and rest were in the middle zone (18%). At 100 lux and 500 lux, percentage of the male wasps sticking to the lower, middle and upper zone did not differ significantly (P>0.05). Only 36% males at 100 lux and 32% at 500 lux were stuck at lower zone whereas, 38% at 100 lux and 40% at 500 lux were stuck to upper zone. At 1000 lux, percentage of wasps that stuck to middle (20%) and upper zone (32%) did not differ significantly whereas, wasps at lower zone (48%) differ significantly with other two zones. Overall, increase in light intensity influenced the flight response of wasps resulting in increased vertical flight of the wasps.

**Fig. 2.**
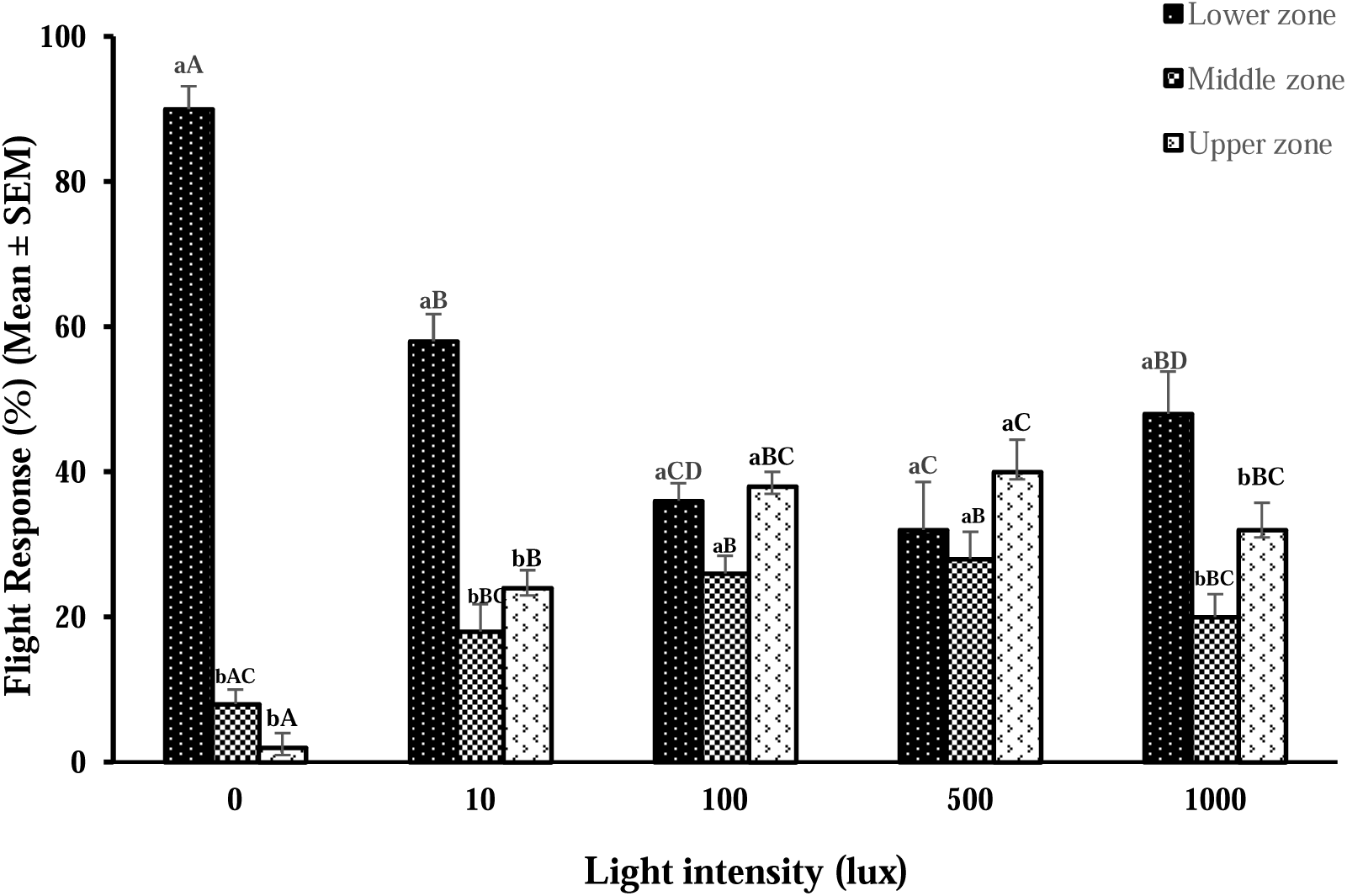
Flight response of male *C. plutellae* wasps at different light intensities in a cylindrical flight chamber. (Bars with different lower case letters differ significantly (p<0.05) on different zone of same light intensity Bars with different upper case letters differ significantly (p<0.05) between different light intensity at particular zone)

The flight response of male wasps was also compared between different light intensities at a particular zone. At lower zone, the percentage of the male wasps that stuck (90%) was significantly higher (P<0.05) at 0 lux than at 10 lux, 100 lux, 500 lux and 1000 lux. The percentage of wasps found sticking on middle zone at 0 lux,10 lux, and 1000 lux and at 100 lux and 500 lux did not differ significantly (P>0.05). At upper zone, the percentage of wasps stuck at 0 lux differed significantly with 10, 100, 500 and 1000 lux. However, the percentage of wasps found sticking to upper zone at 100, 500 and 1000 lux did not differ significantly (P>0.05) (Fig. 2).

When compared between the sexes, it was found that the females were more sensitive to light intensity than males. Both the sexes did not show any significant difference (P>0.05) in the flight activity at 0 lux (Table 1). At 10 lux, significantly higher (P<0.05) percentage of male wasps stuck to the lower zone while females stuck to the upper zone (P<0.001) (Table 1). However, the wasps stuck to the middle zone were statistically similar (P>0.05). At 100 lux, no significant difference (P>0.05) was observed between the sexes stuck on middle zone. Whereas, significantly higher percentage (P<0.001) of male wasps were found stuck to the lower zone and female wasps were found to stick on the upper zone. At 500 and 1000 lux, significantly higher (P<0.001) percentage of female wasps stuck to the upper zone while male wasps were restricted to lower and middle zone (Table 1).

**Table 1.** Flight responseof male and female *C. plutellae* adults at different light intensities in the flight chamber. The response of male and female adults at *p<0.05, **p<0.01, ***p<0.001were significant and ns showed no significance (P = 0.05) (Student’s t-test)

| Light intensity | Zones | Response of wasps |  |  |
| --- | --- | --- | --- | --- |
|  |  | Male | Female | t- value |
| **0 lux** | Lower | 90 ± 3.16 | 78 ± 5.83 | 1.81<sup>ns</sup> |
|  | Middle | 8 ± 2.0 | 12 ± 5.83 | -0.65<sup>ns</sup> |
|  | Upper | 2 ± 2.0 | 10 ± 3.16 | -2.14<sup>ns</sup> |
| **10 lux** | Lower | 58 ± 3.74 | 42 ± 5.83 | 2.31<sup>*</sup> |
|  | Middle | 18 ± 3.74 | 14 ± 8.94 | 0.73<sup>ns</sup> |
|  | Upper | 24 ± 2.45 | 44 ± 5.10 | -3.54<sup>***</sup> |
| **100 lux** | Lower | 36 ± 2.45 | 16 ± 4.0 | 4.26<sup>***</sup> |
|  | Middle | 26 ± 2.45 | 24 ± 2.45 | 0.58<sup>ns</sup> |
|  | Upper | 38 ± 2.0 | 60 ± 4.47 | -4.50<sup>***</sup> |
| **500 lux** | Lower | 32 ± 6.63 | 0 ± 0.0 | 4.82<sup>***</sup> |
|  | Middle | 28 ± 3.74 | 8 ± 2.0 | 4.71<sup>***</sup> |
|  | Upper | 40 ± 4.47 | 92 ± 2.0 | -10.61<sup>***</sup> |
| **1000 lux** | Lower | 48 ± 5.83 | 0 ± 0.0 | 8.23<sup>***</sup> |
|  | Middle | 20 ± 3.16 | 6 ± 2.45 | 3.50<sup>***</sup> |
|  | Upper | 32 ± 3.74 | 94 ± 2.45 | -13.86<sup>***</sup> |

### Flight Response at Different Physiological Condition

The adult *C. plutellae* wasps showed different propensity to initiate the flight at different physiological condition. In control, where freshly emerged naïve female wasps were not provided any food or odour, significantly higher (p<0.05) percentage of wasps (60%) reached to the upper zone of the chamber. Whereas, the percentage of wasps that reached to lower zone (16%) or middle zone (24%) did not differ significantly (Fig. 5.4). When freshly emerged unfed wasps were provided honey volatile from the bottom of the chamber, flight was arrested and significantly higher (p<0.05) percentage of wasps were restricted to lower zone (60%). They either continued to search and antennate over nylon mesh through which honey odour was emanating or they remained stuck to the lower zone (Fig. 3). However, the percentage of wasps that reached to middle zone (14%) was significantly lower (p<0.05) than upper zone (26%) and lower zone (60%) of the chamber.

**Fig.3.**
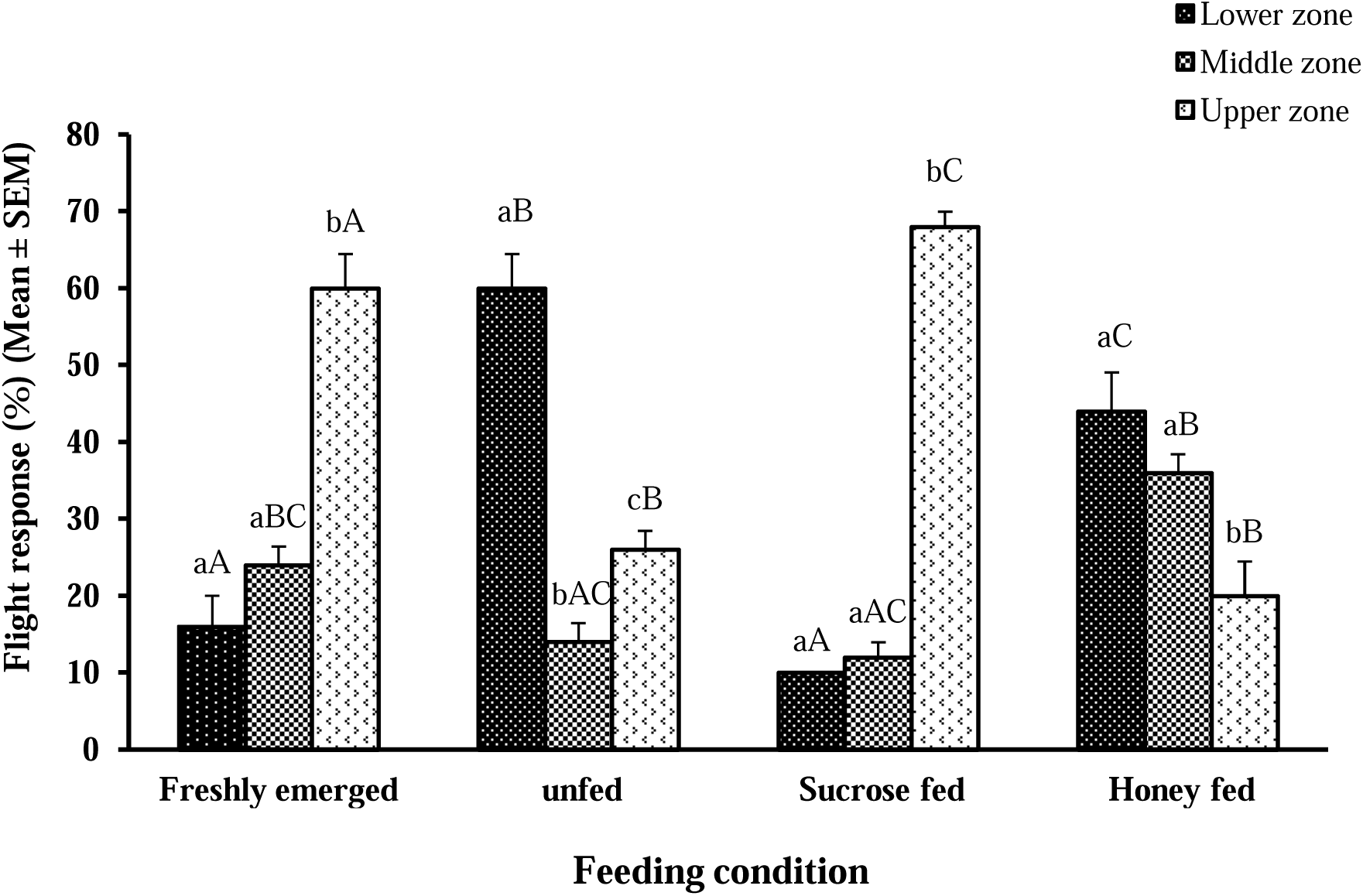
Flight response of female *C. plutellae*wasps in different feeding states in acylindrical flight chamber. (Bars with different lower case letters differ significantly (p<0.05) on different zone of flightchamber at same feeding state. Bars with different upper case letters differ significantly (p<0.05) between different feeding state at particular zone of flight chamber)

When freshly emerged female wasps were fed honey and then provided honey volatile from the bottom of the chamber, odour did not interfere with the flight activity. The percentage of wasps that reached to the upper zone (78%) of the chamber was significantly higher than the other two zones (P<0.05). However, the percentage of female wasps that reached to the lower (10%) and middle zone (12%) did not differ significantly (P>0.05) (Fig. 3). Female wasps fed on sucrose diet did not satiate wasps and their flight activity was arrested by honey odour provided from the bottom of the chamber. Significantly higher (P<0.05) percentage of wasps was observed restricted to the lower zone (44%). They either searched and antennated over nylon mesh or remained stuck to the lower zone as in the case of unfed wasps. The wasps remaining to lower zone (44%) and middle zone (36%) did not differ significantly (P>0.05). However, significantly (P<0.05) lower percentage of wasps reached to upper zone (20%).

The flight activity of the female wasps was also compared between various feeding conditions at a particular zone. At lower zone, unfed female wasps were restricted significantly higher than rest of the other feeding states. However, wasps fed on honey or control did not differ significantly (p>0.05) (Fig. 3). Moreover, at the upper zone, a significantly higher (p<0.05) percentage of female wasps fed on honey was observed than in the wasps at the other feeding state.

The flight activity of male wasps was also influenced by the honey odour. In control, when freshly emerged naïve wasps were not provided any food and honey odour, male wasps reaching to lower (36%), middle (26%) and upper (38%) zones were statistically similar (P>0.05) (Fig. 4). When freshly emerged unfed wasps were provided honey odour from the bottom of the chamber, honey odour arrested the flight activity of male wasps. Significantly (P<0.05) higher percentage of male wasps remained in lower zone (68%). The searching behaviour shown by male wasps were similar to the female wasps. However, wasps that reached to upper zone (26%) was significantly (P<0.05) higher than middle zone (6%) of the chamber (Fig. 4).

**Fig. 4.**
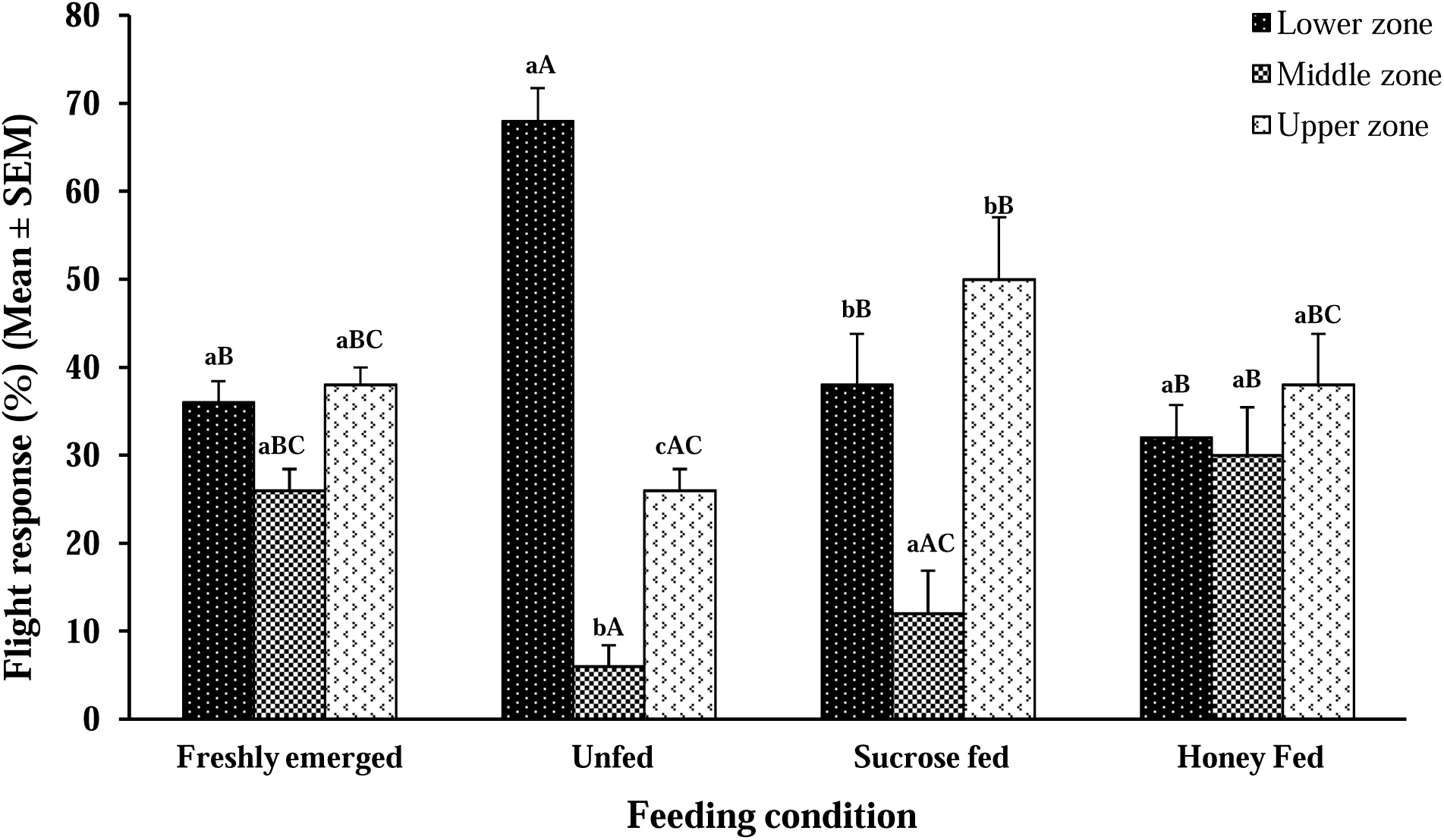
Flight response of male *C. plutellae* wasps, in different feeding state in cylindrical flight chamber. (Bars with different lower case letters differ significantly (p<0.05) on different zone of flight chamber at same feeding state. Bars with different upper case letters differ significantly (p<0.05) between different feeding state at particular zone of flight chamber)

When honey fed male wasps were provided honey volatile, wasps reached to lower (38%) and upper (50%) zone did not differ significantly (P>0.05). Whereas, the wasps reached to middle zone (12%) was significantly (P<0.05) lower than other zones of the chamber. Unlike female wasps, percentage of sucrose fed males reaching to lower (32%), middle (30%) and upper (38%) zone were statistically similar (P>0.05).

Males were also compared for particular zone at various feeding state. The result showed that significantly higher (P<0.05) percentage of unfed wasps remained on lower zone as compared to the rest of the feeding state in this zone. At middle zone, unfed wasps (6%) were stuck significantly (P<0.05) lower than control (26%) and sucrose fed (30%) males. Whereas, at upper zone, honey fed (50%) wasps were significantly higher than rest of the feeding state (Fig. 4).

When the flight activity of both the sexes was compared at each feeding physiological state, it was found that state of feeding significantly influenced the activity of the wasps, especially in female wasps (Table 2). In control, male showed significantly (P<0.001) lower vertical flight (36%) than females (16%) as a result higher percentage of the wasps stuck to lower zone in the absence of any odour (Table 2). Percentage of female wasps that stuck at upper zone (60%) was significantly higher than males (38%). However, there was no significant difference (P>0.05) between male and female wasps on middle zone (Table 2). In the unfed state, no significant difference (p>0.05) was observed in the lower as well as upper zone between the male and female wasps. However, higher percentage of female wasps reached to middle zone as compared to male wasps at unfed state. Statistically lower percentage (P<0.001) of honey fed males stuck to the upper zone (50%) than honey fed females (78%) (Table 2). Higher percentage of the male wasps was restricted to lower zone (38%) than females (10%) in honey fed state. Whereas, there was no significant difference (P>0.05) between male and female at middle zone. In the lower and middle zones, no significant difference (p>0.05) was observed between the sucrose fed males and females. However, significantly higher percentage (p<0.05) of sucrose fed male wasps (38%) was observed in the upper zone than sucrose fed female wasps (20%) (Table 2).

**Table 2.** Flight response of male and female *C. plutellae* adults in different feeding states, in a cylindrical flight chamber. The response of male and female adult at *p<0.05, **p<0.01, ***p<0.001 were significant and ns showed no significance (P = 0.05) (Student’s t-test)

| Feeding condition | Zones | Response of wasps |  |  |
| --- | --- | --- | --- | --- |
|  |  | Male | Female | t-value |
| Freshly emerged | Lower | 36 ± 2.45 | 16 ± 4.0 | 4.26<sup>***</sup> |
|  | Middle | 26 ± 2.45 | 24 ± 2.45 | 0.58<sup>ns</sup> |
|  | Upper | 38 ± 2.0 | 60 ± 4.47 | -4.49<sup>***</sup> |
| Without fed | Lower | 68 ± 3.74 | 60 ± 4.47 | 1.37<sup>ns</sup> |
|  | Middle | 6.0 ± 2.45 | 14 ± 2.45 | -2.31<sup>*</sup> |
|  | Upper | 26 ± 2.45 | 26 ± 2.45 | 0.00<sup>ns</sup> |
| Honey fed | Lower | 38 ± 5.83 | 10 ± 0.0 | 4.80<sup>***</sup> |
|  | Middle | 12 ± 4.90 | 12 ± 2.0 | 0.00<sup>ns</sup> |
|  | Upper | 50 ± 7.07 | 78 ± 2.0 | -3.81<sup>***</sup> |
| Sucrose fed | Lower | 32 ± 3.74 | 44 ± 5.10 | -1.90<sup>ns</sup> |
|  | Middle | 30 ± 5.48 | 36 ± 2.45 | -1.00<sup>ns</sup> |
|  | Upper | 38 ± 5.83 | 20 ± 4.47 | 2.45<sup>*</sup> |

## Discussion

Flight allows insect parasitoid to disperse over long distances in search for food, partner and oviposition sites across a wide range of environment. In the present study, the flight activity was measured in terms of vertical angle of flight in parasitoids which in turn help the parasitoid to take high flight and disperse from the releasing sites. In this study, both the sexes remain inactive in the dark (0 lux light intensity) and do not show any vertical angle of flight. As the light intensity increases, vertical angle of flight also increases reflected by percentage of wasps reaching and sticking to the upper zone in the flight chamber. However, females showed high response at higher light intensity than males. This indicates that the females are more active flier than the males. Fahrner et al (2014) also reported that females of *Tetrastichus planipennisi* dispersed much farther than male parasitoids. The adults of *Eretmocerus eremicus* wasps were also able to sustain direct flight towards the sky light cues (Blackmer and Cross 2001). Another study on *C. glomerata* showed that cloudy and weather may suppress the flight activity and higher light intensity increase this activity (Gu and Dorn 2001). Barbosa and Frongillo (1977) also observed that flight and locomotory activity of *Brachymeria intermedia* increased as the light intensity increased and activity was dramatically depressed in the total darkness. Adults of *Tirumala limniace* also exhibited increased flight activity and taking flight earlier at high intensity of light as compared to weak light (Liao et al. 2017).

The gender related differences in resource allocation are commonly found in parasitoids (Rivero and West 2002; Wanner et al. 2006). In most of the cases, females always show more active response to a given cues than males like flight behaviour (Bellamy and Byrne 2001). In present study, both sexes of *C. plutellae* exhibited varied level of vertical flight under similar light condition in the flight chamber. Higher percentage of female *C. plutellae* showed higher vertical flight than males with increase in light intensity. Females of *Eretmocerus eremicus* showed 10.6 times longer flight duration and more responsive to land on plant cues as compare to males (Bellamy and Byrne 2001). The females have to fly over larger distance to search host while male wasps normally stay close to natal patch, mating with emerging females (Laing and Levin 1982; Wanner et al. 2006).

The physiological state of parasitoid can also affect their flight activity. Honey odour, which was provided from bottom of flight chamber, was able to arrest flight activity of unfed *C. plutellae,* and wasps were continuously antennating and searching for honey over nylon net and try to reach the food source. It indicates that the starved adults are not efficient in flight and also parasitising the host as they are busy in food searching. In contrast, honey fed females did not show any arrestment behaviour by the odour and majority of females reached to the upper zone. Same pattern of flight response was observed in adult male wasps. The distance of fight in adult parasitoid, *Tetrastichus planipennisi* fed honey mixed in water, was longer as compare to those which were fed only water, indicating that energy acquired during larval stage is not suffcient to fuel adult fight (Fahrner et al 2014). In fact, it appears necessary to provide *T. planipennisi* a honey solution before it releases into the field to achieve maximum fight.

Energetically, flight is most costly activity of parasitoids (Casas et al. 2003; Hoferer et al. 2000), and unfed parasitoids exhaust their energy reserves at considerably faster rates under field condition as compared to individuals kept in confined conditions (Steppuhn and Wäckers 2004). Rousse et al (2009) observed that two days of starvation reduced the fight activity of female *Fopius arisanus* and on third day, they started responding to food (honey) rather than host stimuli (host feces). They also concluded that carbohydrate shortage in the field limits the fight induction and therefore infuenced by the accessibility of carbohydrate sources (such as honeydew or foral nectar). In *Cotesia glomerata,* a synovigenic parasitoid species, fed adult female wasps are able to increase their flight activity (Wanner et al. 2006).

*C. plutellae* adults fed on sucrose, in the present study, did not show efficient flight behaviour as in case of honey fed adults. Also, sucrose fed wasp did not initiate take off in the presence of honey odour. It might be possible that the sucrose feeding does not provide the entire nutritional requirement to the wasps. Presence of honey odour elicited innate response from adult wasps that continuously search and antennate on honey odour, and as a result, wasps showed poor flight response.

It has already been shown that the honey has larger effect on the flight parameter of *C. glomerata* than a single sugar component like sucrose and it appears to be less suitable for sustaining long flight activity (Wanner et al. 2006). Though, carbohydrates are known to be predominant substrate for flight in most of the Hymenoptera and Diptera (Beenakker et al. 1984). The importance of food odour in attracting various beetles has also been demonstrated by Barrer (1983). Wanner et al. (2006) showed that nectars with different nutritional components exhibit different effect on flight capacity of parasitoid wasps, *Cotesia glomerata* (L.) (Hymenoptera: Braconidae). In contrast to these studies, Fischbein et al. (2011), showed that the different flight parameters in *Ibalia leucospoides* (H) (Hymenoptera: Ibaliidae) are not affected by prior access to food source and such effect may manifest itself on subsequent days of parasitoid flight. Therefore, it is widely accepted that carbohydrate rich food supplements are important for parasitoids prior to their release in the field for enhancing paraitisation and regulation of pest population in the field.

The present study suggests that feeding of parasitoid on food source plays a crucial role in success of biological control program. The results of these experiments provide valuable information for determining the suitable food source which enhances flight activity and alternatively increase the searching efficiency of female parasitoid wasps to search host insects in larger field area which ultimately increase parasitisation and boost the biological control program. However, further research is needed to provide experimental evidence of the dynamics of nutrients utilization during flight and the potential role of that food on other life history traits (longevity and fecundity and other physiological parameters) of *C. plutellae.*

## Acknowledgements

We thank University of Delhi, India, for providing facilities in the Department of Zoology. We gratefully acknowledge Research Fellowship provided by the University Grants Commission for this work.

## References

Alves TJS, Silva-Torres CSA, Wanderley-Teixeira V, Teixeira AAC, Torres JB, Lima TA, Ramalho FS (2015). Behavioral studies of the aprasitoid *Bracon vulgaris* Ashmead (Hymenoptera: Braconidae). J Insect Behav 28(5):604–617.

Andrewartha HG, Birch LC (1954) “The Distribution and Abundance of Animals.” Univ. of Chicago Press, Chicago.

Barbosa P, Frongillo EA Jr. (1977) Flight and locomotory responses of *Brachymeria intermedia* (Nees) [Hym.: Chalcididae] in various temperatures and light intensities. Entomoph 22:405–411.

Barrer PM (1983) A field demonstration of odour-based, host-food finding behaviour in several species of stored grain insects. J Stored Prod Res 19:105–110.

Beenakkers DJ, Horst VD, Van Merrievic WJA (1984) Insect flight muscle metabolism. Insect Biochem 14(3):243–260.

Bellamy DE, Byrne DN (2001) Effects of gender and mating status on self directed dispersal by the whitefly parasitoid *Eretmocerus eremicus*. Ecol Entomol 26:571–577.

Blackmer JL, Cross D (2001) Response of *Eretmocerus eremicus* to skylight and plant cues in a vertical flight chamber. Ent Exp Appl 100:295–300.

Casas J, Driessen G, Mandon N, Wielaard S, Desouhant E, van Alphen JJM, Lapchin L, Rivero A, Christides JP, Bernstein C (2003) Energy dynamics in a parasitoid foraging in the wild. J Ani Ecol 72:691–697.

Chapman RF (1998) The insects: structure and function. Cambridge Univiversity Press, Cambridge.

Eijs I, Ellers J, van Duinen G (1998) Feeding strategies in drosophilid parasitoids: the impact of natural food resources on energy reserves in females. Ecol Entomol 23:133–138.

Fahrner SJ, Lelito JP, Blaedow K, Heimpel GE, Aukema BH (2014) Factors affecting the flight capacity of *Tetrastichus planipennisi* (Hymenoptera: Eulophidae), a classical biological control agent of *Agrilus planipennis* (Coleoptera: Buprestidae). Environ Entomol 43:1603–1612.

Fischbein D, Corley JC, Villacide JM, Bernstein C (2011) The influence of food and con-specifics on the flight potential of the parasitoid *Ibalia leucospoides*. J Insect Behav. 24: 456–467.

Gaudon JM, Allison JD, Smith SM (2018) Factors influencing the dispersal of a native parasitoid, *Phasgonophora sulcata*, attacking the emerald ash borer: implications for biological control. BioCont DOI: 10.1007/s10526-018-9900-x

Gaudon JM, Haavik LJ, MacQuarrie CJK, Smith SM, Allison JD (2016) Influence of nematode parasitism, body size, temperature, and diel periodicity on the flight capacity of *Sirex noctilio* F. (Hymenoptera: Siricidae). J Insect Behav 29:301–314.

Gu H, Dorn S (2001) How do wind velocity and light intensity influence host location success in *Cotesia glomerata* (Hym., Braconidae). J Appl Entomol 125:115–120.

Hoferer S, Wäckers FL, Dorn S (2000) Measuring CO_2_ respiration rates in the parasitoid *Cotesia glomerata*. Mitteilungen der DeutschenGesellschaft fuer allgemeine undangewandte Entomol 12:555–558.

Idris AB, Grafius E (1995) Wild flowers as nectar sources for *Diadegma insulare* (Hymenoptera: Ichneumonidae), a parasitoid of diamondback moth (Lepidoptera: Yponomeutidae). Environ Entomol 24:1726–1735.

Jerbi-Elayed M, Lebdi-Grissa K, Goff GL, Hance T (2015) Influence of temperature on flight, walking and oviposition capacities of two aphid parasitoid species (Hymenoptera: Aphidiinae). J Insect Behav 28(2):157–166.

Jervis MA, Kidd NAC, Fitton MG, Huddleston T, Dawah HA (1993) Flower visiting by hymenopteran parasitoids. J Nat History 27:67–105.

Keppner EM, Jarau S (2016) Influence of climatic factors on the flight activity of the stingless bee Partamona orizabaensis and its competition behavior at food sources. J. Comp. Physiol. 202: 691–699.

Laing JE, Levin DB (1982) A review of the biology and a bibliography of *Apanteles glomeratus* (L.) (Hymenoptera: Braconidae). Bioc News Inf 3:7–23.

Lewis WJ, Stapel JA, Cortesero AM, Takasu K (1998) Understanding how parasitoids balance food and host needs: importance to biological control. Biol Cont 11:175–183.

Liao H, Shi L, Liu W, Du T, Ma Y, Zhou C, Deng J (2017) Effects of light intensity on the flight behaviour of adult *Tirumala limniace* (Cramer) (Lepidoptera: Nymphalidae). J Insect Behav 30(2):139–154.

Luo L, Johnson SJ, Hammond AM, Lopez JD, Geaghan JP, Beerwinkle KR, Westbrook JK (2002) Determination and consideration of flight potential in a laboratory population of true armyworm (Lepidoptera: Noctuidae). Environ Entomol 31:1–9.

Mills NJ, Heimpel GE (2018) Could increased understanding of foraging behavior help to predict the success of biological control? Curr Opin Insect Sci. 27: 26–31.

Rivero A, West SA (2002) The physiological costs of being small in a parasitic wasp. Evol Ecol Res 4:407–420.

Rousse P, Gourdon F, Roubaud M, Chiroleu F, Quilici S (2009) Biotic and Abiotic Factors Affecting the Flight Activity of *Fopius arisanus*, an Egg-Pupal Parasitoid of Fruit Fly Pests. Environ. Entomol. 38(3): 896–903.

Sirot E, Bernstein C (1996) Time sharing between host searching and food searching in parasitoids: state-dependent optimal strategies. Behav Ecol 7:189–194.

Steppuhn A, Wäckers FL (2004) HPLC sugar analysis reveals the nutritional state and the feeding history of parasitoids. Funct Ecol 18:812–819.

Wäckers FL (2005) Suitability of (Extra-) floral nectar, pollen, and honeydew as insect food sources. In: Wäckers, F.L., van Rijn, P.C.J., Bruin, J. (Eds.), Plantprovided food for carnivorous insects: a protective mutualism and its applications, Cambridge university press, UK, pp. 17–74.

Wanner H, Gu H, Dorn S (2006) Nutrition value of floral nectar sources for flight in the parasitoid wasp, *Cotesia glomerata*. Physiol Entomol 31:127–133.

Yu H, Zhang Y, Wu K, Wyckhuys KAG, Guo Y (2009) Flight potential of *Microplitis mediator*, a parasitoid of various lepidopteran pests. BioCont 54:183–193.

